# Matrix-Bound Growth Factors are Released upon Cartilage Compression by an Aggrecan-Dependent Sodium Flux that is Lost in Osteoarthritis

**DOI:** 10.1101/2021.01.31.428791

**Authors:** Stuart J. Keppie, Jessica C. Mansfield, Xiaodi Tang, Christopher J. Philp, Helen K. Graham, Patrik Önnerfjord, Alanna Wall, Celia McLean, C. Peter Winlove, Michael J. Sherratt, Galina E. Pavlovskaya, Tonia L Vincent

**Author notes:** Correspondence, Kennedy Institute of Rheumatology, Centre for OA Pathogenesis Versus Arthritis, University of Oxford, Roosevelt Drive, OX3 7FY, United Kingdom.

## Abstract

Articular cartilage is a dense extracellular matrix-rich tissue that degrades following chronic mechanical stress, resulting in osteoarthritis (OA). The tissue has low intrinsic repair especially in aged and osteoarthritic joints. Here we describe three pro-regenerative factors; fibroblast growth factor 2 (FGF2), connective tissue growth factor, bound to transforming growth factor-beta (CTGF-TGFβ), and hepatoma-derived growth factor (HDGF), that are rapidly released from the pericellular matrix (PCM) of articular cartilage upon mechanical injury. All three growth factors bound heparan sulfate, and were displaced by exogenous NaCl. We hypothesised that sodium, sequestered within the aggrecan-rich matrix, was freed by injurious compression, thereby enhancing the bioavailability of pericellular growth factors. Indeed, growth factor release was abrogated when cartilage aggrecan was depleted by IL-1 treatment, and in severely damaged human osteoarthritic cartilage. A flux in free matrix sodium upon mechanical compression of cartilage was visualised by ^23^Na magnetic resonance imaging (MRI) just below the articular surface. This corresponded to a region of reduced tissue stiffness, measured by scanning acoustic microscopy and second harmonic generation microscopy, and where Smad2/3 was phosphorylated upon cyclic compression. Our results describe a novel intrinsic repair mechanism, controlled by matrix stiffness and mediated by the free sodium concentration, in which heparan sulfate-bound growth factors are released from cartilage upon injurious load. They identify aggrecan as a depot for sequestered sodium, explaining why osteoarthritic tissue loses its ability to repair. Treatments that restore matrix sodium to allow appropriate release of growth factors upon load are predicted to enable intrinsic cartilage repair in osteoarthritis.

**Significance Statement:** Osteoarthritis is the most prevalent musculoskeletal disease, affecting 250 million people worldwide ^1^. We identify a novel intrinsic repair response in cartilage, mediated by aggrecan-dependent sodium flux, and dependent upon matrix stiffness, which results in the release of a cocktail of pro-regenerative growth factors after injury. Loss of aggrecan in late-stage osteoarthritis prevents growth factor release and likely contributes to disease progression. Treatments that restore matrix sodium in osteoarthritis may recover the intrinsic repair response to improve disease outcome.

## Introduction

Articular cartilage is a highly mechanosensitive tissue that overlies the ends of bone in all articulating joints. It is predominantly composed of extracellular matrix, with the remaining tissue containing sparsely distributed cells (chondrocytes) that constitute 5—10% of the tissue by volume. Articular cartilage contains an abundance of type II collagen, a fibrillar collagen responsible for providing tensile strength to the tissue, and proteoglycans. The most abundant proteoglycan of articular cartilage is aggrecan, to whose core protein are attached ~100 chondroitin sulfate glyscosaminoglycan (GAG) chains ^2^, which generate a net negative fixed charge to the tissue. This negative charge contributes to the osmotic retention of water in the tissue by attracting sodium cations, and provides intermolecular electrostatic repulsion between GAGs, contributing to tissue stiffness and elasticity. The aggrecan-rich extracellular matrix is divided into the territorial matrix (TM) and interterritorial matrix (ITM) and these are separated from the cell by a distinct region of matrix termed the pericellular matrix (PCM), which is rich in collagen VI and the heparan sulfate proteoglycan perlecan ^3^.

Loss of aggrecan is an early pathological feature of cartilage matrix breakdown that occurs in osteoarthritis (OA), a disease estimated to affect over 250 million people worldwide ^4^, and a leading cause of disability and societal burden. Other disease features of the OA joint include synovial hypertrophy, osteophyte formation and subchondral bone sclerosis. Degradation of aggrecan can be detected histologically using cationic dyes, as well as in patients *in vivo* by sodium-MRI imaging ^5^. A number of pathogenic mechanisms contribute to the development of OA, which can be broadly divided into those that drive proteolytic degradation of the matrix and those that suppress the intrinsic repair of the tissue. Both are closely linked to the two major aetiological factors in disease; abnormal mechanical joint loading and ageing.

In exploring the response of articular cartilage to mechanical stress, our group has observed that sequestered matrix-bound growth factors are released from the pericellular matrix of cartilage following mechanical injury. The best characterised of these is fibroblast growth factor 2 (FGF2), which is chondroprotective *in vivo* in multiple animal models of OA ^6–12^. The second molecule identified was connective tissue growth factor (CTGF), which is in a covalently bound complex with latent transforming growth factor beta (TGFβ) ^13^. When the latent complex is released from cartilage upon injury, latent TGFβ is activated by binding to the cell surface receptor TGFβR3 ^13^. TGFβ is a strong chondrogenic growth factor ^14,15^ and is implicated in intrinsic tissue repair responses elsewhere in the body ^16-18^. TGFβ and FGF2 exhibit binding affinities to heparin and heparan sulfate ^19^. In our previous studies both growth factors were released rapidly upon injury (within 5 minutes), and this was independent of cell viability ^13,20^.

In this study, we characterise a third growth factor released upon cartilage injury, hepatoma-derived growth factor (HDGF). We reveal the common mechanism by which these growth factors are released upon tissue injury and how this goes awry in OA.

## Results

### PCM Growth Factors, Including Hepatoma Derived Growth Factor (HDGF) are Bound to Heparan Sulfate and Released Rapidly upon Cartilage Injury

We previously identified HDGF from a proteomic analysis of human isolated PCM and tested whether HDGF was released upon cartilage injury in a similar fashion to release of FGF2 and CTGF. FGF2, CTGF and HDGF were all released from human knee cartilage within 30 minutes of recutting injury (Figure 1A), and were not detected in the medium of uninjured, rested cartilage explants. HDGF release into the medium occurred largely within the first 5 minutes (Figure 1B). We confirmed our previous observation of co-localisation of FGF2 and CTGF in the type VI collagen and perlecan-rich pericellular matrix (Figure 1C) and confirmed heparin binding of all three growth factors by showing their depletion from the injury conditioned medium (recut CM) on a heparin column, and elution with 2.5M NaCl (Figure 1D). They were also released from articular cartilage following treatment with heparitinase (Figure 1E). The injurious release of growth factors occurred irrespective of whether the tissue was alive or dead (Figure 1F), at 4 °C or 37 °C, or after pre-incubation with an inhibitor of matrix metalloproteases, or a broad-spectrum protease inhibitor (Figure 1G). These results suggested that there was a common mechanism of release of PCM-bound growth factors upon cartilage injury, which did not require viable cells or involve cell-dependent protease activity.

**Fig. 1.**
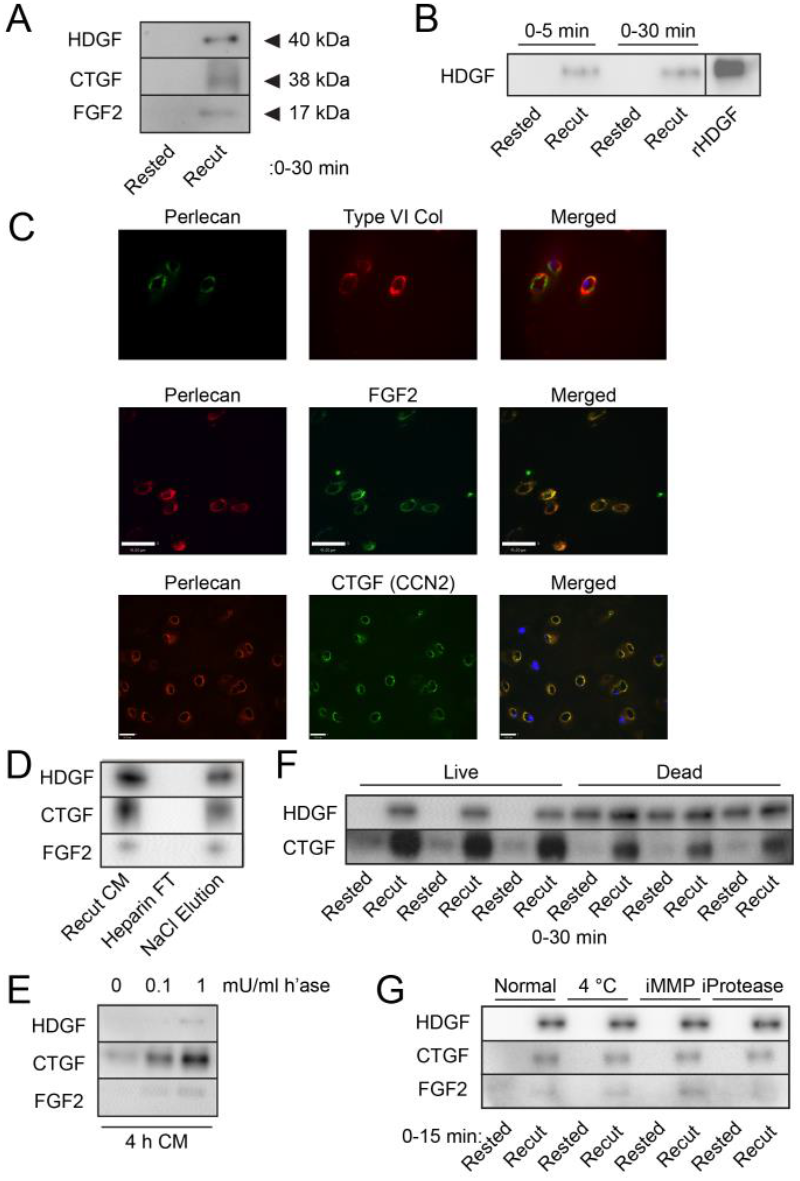
Heparan sulfate-bound growth factors are released upon cartilage injury. (A) Immunoblots of medium conditioned (30 min) by rested (48 h post dissection) or recut (Recut CM) human articular cartilage. (B) Medium conditioned by rested and recut cartilage for 5 or 30 minutes. Recombinant HDGF (rHDGF). (C) confocal microscopy showing co-immunolocalisation of type VI collagen/perlecan, perlecan/FGF2 and perlecan/CTGF (CCN2) in the PCM. (D) Injury CM (recut) was passed through a heparin-agarose column and eluted with 2.5M NaCl. Starting material (Recut CM), column flow through (Heparin FT) and eluted fractions (NaCl Elution) were immunoblotted for CTGF, HDGF and FGF2. (E) Rested porcine articular cartilage was treated with 0, 0.1 or 1 mU/ml heparitinase I for 4 h and the conditioned medium immunoblotted for growth factors. (F) Immunoblots showing release of FGF2, CTGF and HDGF in rested or recut CM (30 mins) from live or dead (3 cycles of freeze-thawing) porcine articular cartilage. (G) Porcine articular cartilage explants were pre-treated with 10 μM batimastat (MMP inhibitor) or with protease inhibitor cocktail, or were incubated at 4°C or 37°C prior to recutting.

### Manipulation of Tissue Sodium Affects Growth Factor Release upon Cartilage Injury

As the interaction of heparin-bound growth factors with heparan sulfate is ionic and the growth factors in injury CM were displaced by elution from a heparin column with sodium chloride (Figure 1D), we hypothesised that growth factor release after injury was dependent upon mobilisation of tissue sodium. This hypothesis was supported by the factthat the PCM was devoid of chondroitin sulfate (Figure 2A), indicating a cation gradient from the territorial to the pericellular matrix.

**Fig. 2.**
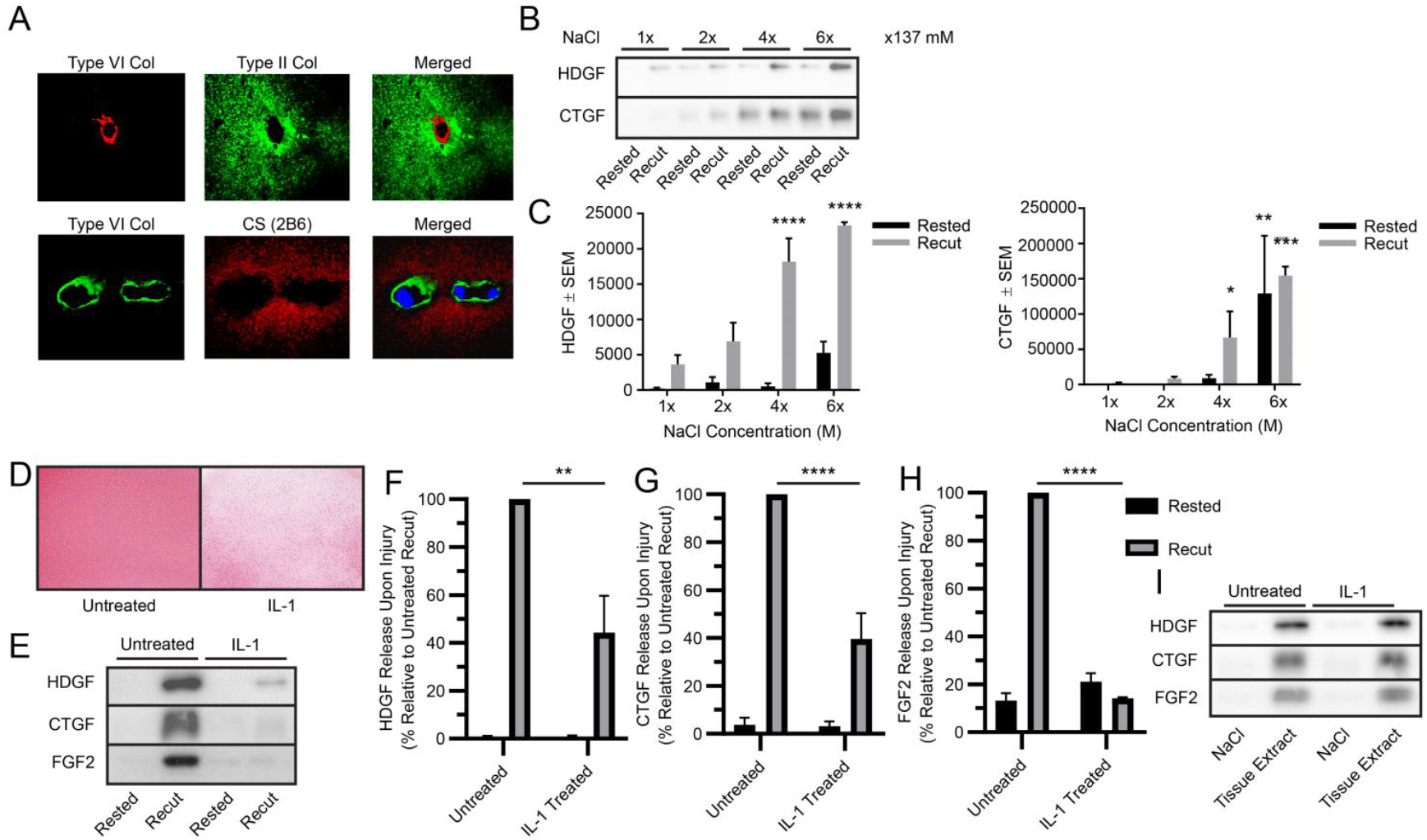
Changes in tissue NaCI affect injury-induced growth factor release. (A) Confocal microscopy delineating pericellular matrix (type VI collagen) and territorial matrix (type II collagen and chondroitin sulfate (CS)), in human articular cartilage. (B) Western blot of CM (0-15 mins) from rested and recut porcine articular cartilage generated in medium with physiological NaCl (0.137 mM) or supraphysiological NaCl concentrations (2-6 fold). (C) Quantification for HDGF and CTGF (Two-Way ANOVA with Sidak multiple comparisons compared to 1x NaCl, n=3 biological replicates). (D-I) Aggrecan was depleted from articular cartilage by 7 day IL-1 treatment. (D) Safranin O staining of porcine cartilage after 7 days. (E) Immunoblots of growth factors of rested and injury CM after recutting (0-15 min). (F-H) Quantification of western blots for HDGF, CTGF and FGF2 respectively (T-test, n=6 biological replicates). (I) Western blots of tissue extracts (2.5M NaCl or dissociative lysis buffer) of control and IL-1-treated explants (representative experiment of four independent experiments). (*=p<0.05, **=p<0.01, ***=p<0.001, ****=p<0.0001).

To test the hypothesis, we studied the injury response after manipulating tissue sodium levels. Addition of exogenous sodium chloride, at supraphysiological concentrations, displaced HDGF and CTGF from cartilage in a concentration dependent manner and enhanced the release of HDGF and CTGF after injury (Figure 2B, C). Conversely, depletion of aggrecan from cartilage following prolonged treatment with interleukin-1 (IL-1), reduced the fixed charge of the tissue (Figure 2D) and abrogated the release of HDGF, CTGF and FGF2 upon recutting (Figure 2E–H). The reduction in release upon injury occurred despite there being no difference in total growth factor extracted from untreated and IL-1 treated tissue (Figure 2I).

### Bound Sodium is Mobilised upon Cartilage Compression

We next examined the effect of cartilage mechanical compression on sodium mobilisation using multi-modal MRI methods (Figure A). Porcine knee osteochondral plugs were imaged using a Bruker AVANCE III 9.4T microimaging system at micron length scales to obtain anatomical, free and bound sodium scans of cartilage plugs during compressive loading. A strong sodium signal was seen in the articular cartilage, but not subchondral bone (Figure 3B). The MRI signal contribution from bound sodium was determined by inserting a multiple quantum filter before applying localising gradients (Supplementary Figure S2). Tissue compression (28% strain) caused redistribution of both free and bound sodium within articular cartilage (Figure 3B). Compression led to a heterogeneous distribution in free sodium in the cartilage matrix, specifically just below the superficial layer. Compression reduced bound sodium levels, with partial restoration of these levels upon relaxation of the cartilage. This was further confirmed by two dimensional ^23^Na TQTPPI spectroscopy (details in supplementary - ^23^Na MRS) where sodium signals in both free and bound states are recorded, providing information about bulk sodium levels in each state. Tissue compression caused a significant increase in free sodium of 46%, with a concomitant significant decrease in the bound sodium fraction (F(2,9) = 9.709, p=0.0057) (Figure 3C). Upon relaxation of the tissue, the free and bound sodium fractions were lower than in the compressed state, with a tendency towards uncompressed levels.

**Fig. 3.**
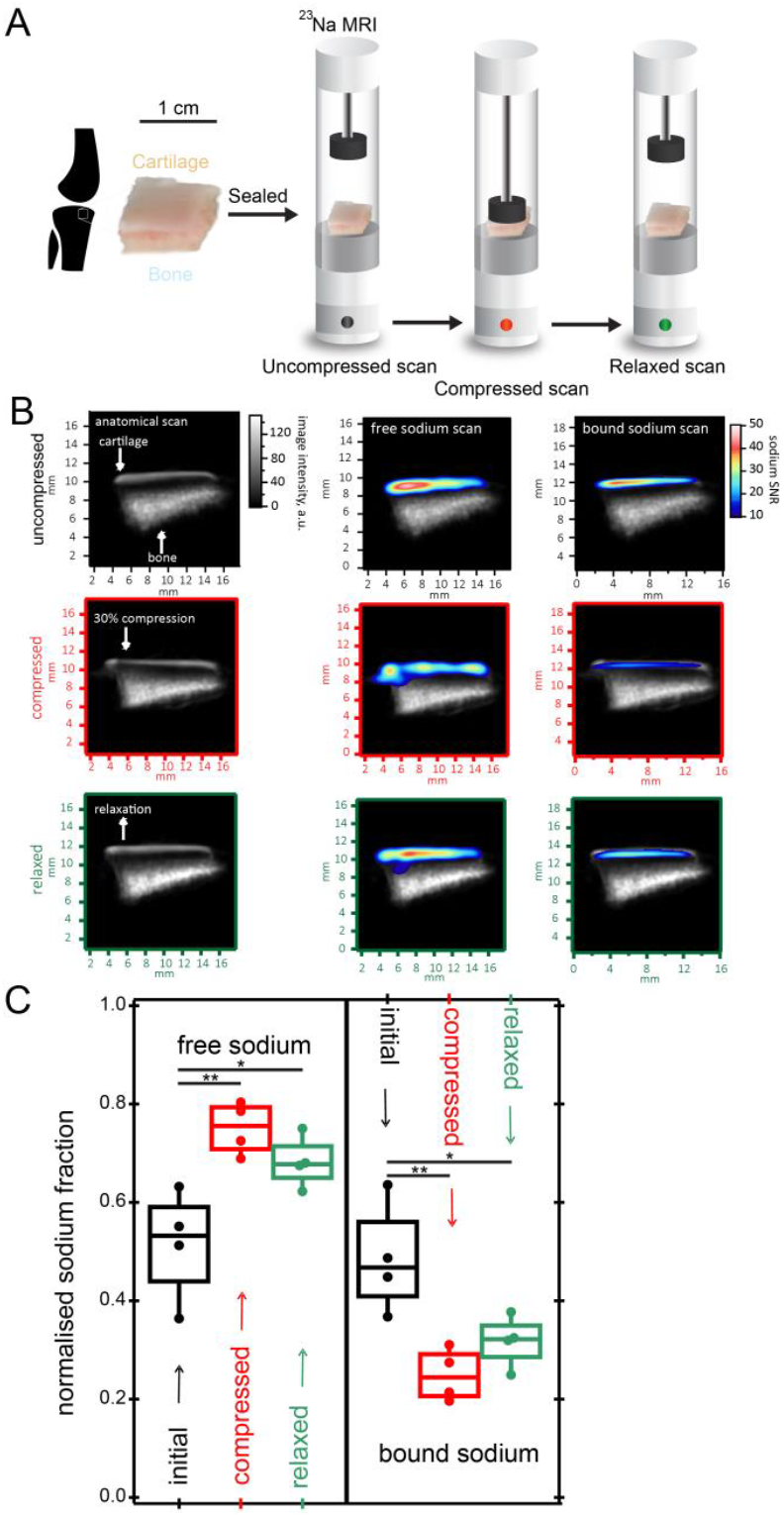
Sodium is reversibly mobilised from bound stores upon cartilage compression. (A) Schematic of tibial plateau osteochondral plugs for ^1^H (anatomical) and ^23^Na (free and bound) MRI performed at microscale resolution in initial (uncompressed), compressed (28% strain) and relaxed states. All images were recorded at steady state. (B) Anatomical, free and bound sodium scans overlaid with corresponding anatomical scans of vacuum-sealed porcine osteochondral plugs during loading cycle depicted in A. (C) Free and bound sodium fractions (n=4) quantified with 2D TQTPPI ^23^Na spectroscopy (details in supplementary) in the initial, compressed and relaxed states as depicted in A. One-Way ANOVA with Tukey multiple comparisons (*=p<0.05, **=p<0.01).

### Compression-Induced Growth Factor Release is Preferentially in Areas of Low Matrix Stiffness

To explore further the possibility that regional differences exist within the tissue, cyclically compressed porcine knee cartilage was examined by immunohistochemistry for evidence of growth factor dependent cell activation. SMAD2 phosphorylation, indicative of TGFβ activation, was evident in the superficial zone of cartilage from just below the articular surface (between 50 and 300 μm from the surface) in compressed, but not non-compressed cartilage explants (Figure 4A–C). Regional activation was not due to differences in the amount of growth factor present in the tissue at different levels, as all three PCM growth factors were identified throughout the deeper parts of the tissue by proteomic analysis (Figure 4D). Full proteomic dataset of each cartilage layer shown in supplementary file and raw data available via ProteomeXchange with identifier PXD023890.

**Fig. 4.**
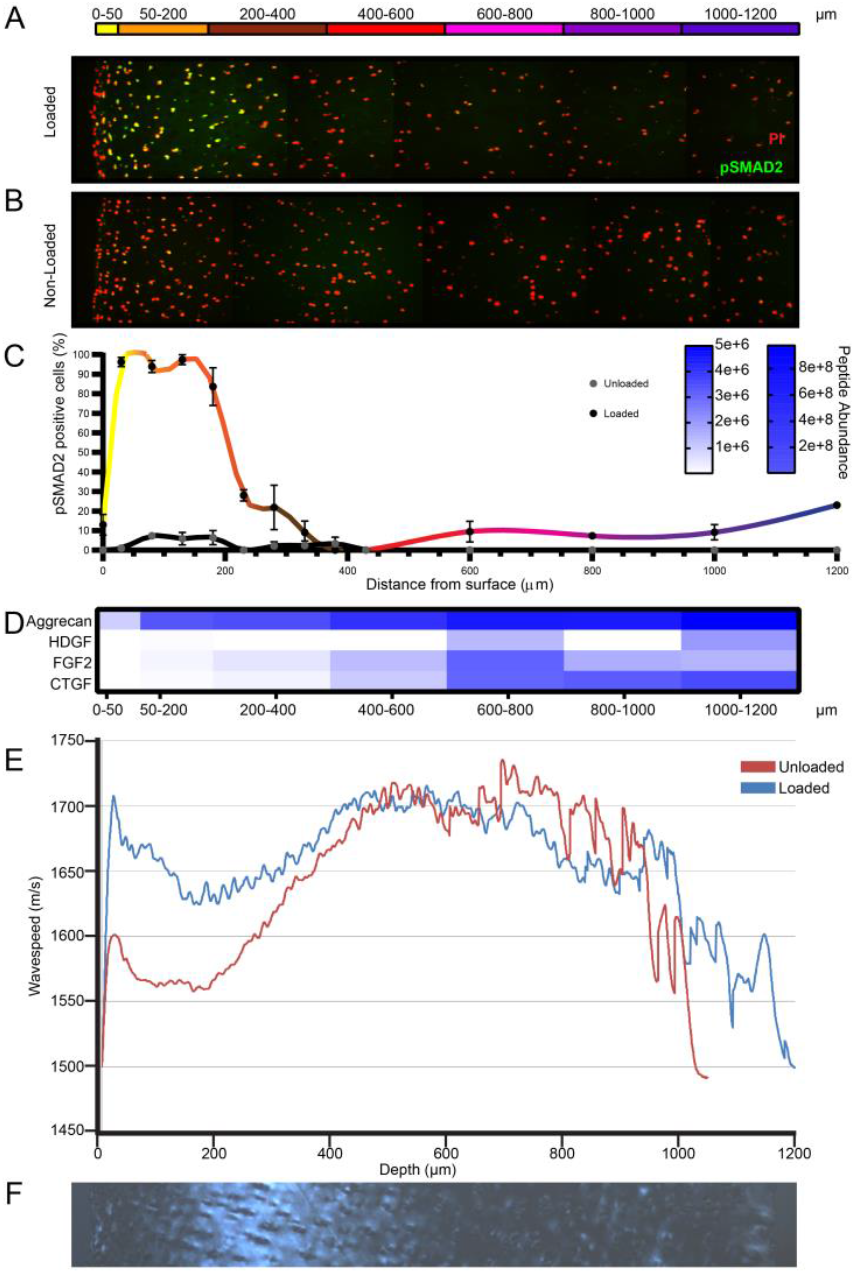
TGFβ/CTGF mediated SMAD2 activation occurs after cyclical compression in a region of cartilage with low tissue stiffness. Porcine articular cartilage was (A) cyclically loaded for 45 min or (B) left unloaded. (A,B) Coronal sections perpendicular to the articular surface were immunostained for phosphorylated SMAD2 (green) and (C) quantified (n=6 separate experiments). Nuclei were counterstained with propidium iodide. (D) Slices of articular cartilage, parallel with the articular surface were removed by cryosection and analysed by proteomics. (E) Scanning acoustic microscopy (SAM) was performed across the depth of porcine knee cartilage sections, where wavespeed denotes tissue stiffness from the surface (left) to the deep zone (right). (F) Articular cartilage sections were imaged for collagen alignment by polarised light microscopy.

As the region of SMAD2 activation was not apparently limited by growth factor distribution within the tissue, we next explored whether regional differences in tissue stiffness might explain differential activation upon compression. Scanning acoustic microscopy wave speeds were recorded across the depth of the tissue in compressed and non-compressed tissue sections. Variations in stiffness through the depth of the tissue were observed, with the highest stiffness (highest wave speed) in the most superficial 50 μm and a region of reduced stiffness below this from 50– 350μm, which became more evident upon compression (Figure 4E). This region also contained a higher polarised light microscopy signal, consistent with disordered collagen fibres in this region (Figure 4F).

Differential stiffness and response to compression was further investigated in human articular cartilage by second harmonic generation, to explore the collagen organisation and arrangement upon compression. Following compressive load of the tissue (0–21% strain), a marked loss of tissue volume was observed in the superficial zone (Figure 5A, upper box), compressive load led to marked loss of tissue volume (Figure 5B-D), with reorganisation of the collagen fibre orientation (Figure 5E). In contrast, compression of the tissue led to no discernible volume loss in the deep region (Figure 5A lower box, Figure 5F-H), with no change in the collagen orientation (Figure 5I). In both regions of interest, the submicron organisation parameter *I*_2_ which represents the degree of order of individual collagen fibrils within a focal spot, remained constant (data not shown). Taken together, these data confirm region-specific difference in tissue stiffness and response to compression in human cartilage, similar to that seen in pig.

**Fig. 5.**
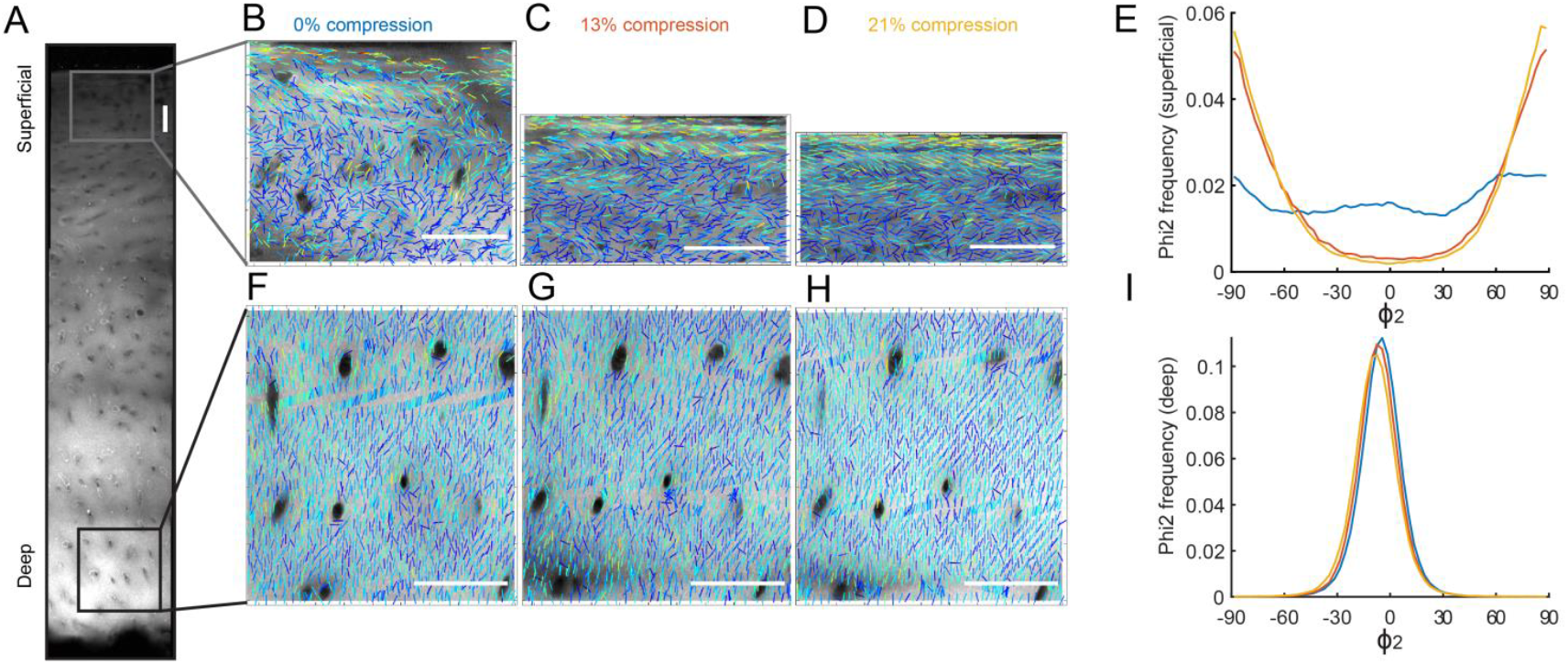
Polarisation sensitive second harmonic generation (SHG) measurement of collagen organisation of human articular cartilage under compressive load. (A) Full depth image of the unloaded region of cartilage, highlighting the 2 areas where polarisation SHG analysis was carried out at 0% (B, F), 13% (C, G) and 12% (D, H). These correspond to the unloaded samples and then the overall strain in the full thickness of the cartilage. Maps of the collagen organisation in the (B-D) superficial region and (F-H) deep region, as a function of load. The direction of the sticks indicates the collagen fibre direction and the colour of the sticks indicates *I*_2_ (level of submicron organisation). φ_2_ measurements, indicating a change in orientation of collagen fibres upon compression, are shown in the (E) superficial layer and the (I) deep layer. Blue line (0% stain), orange line (13% strain), yellow line (21% strain). Note the marked change in compression and collagen orientation in the superficial, but not deep, region. Scale 100 μm.

### Severely Damaged Osteoarthritic Tissue Exhibits Impaired Release of Heparan Sulfate Bound Growth Factors after Injury

Proteolytic degradation of aggrecan in OA occurs through the actions of aggrecanases ^21–23^ and results in a reduction of the fixed charge density of the tissue reflected by the loss of safranin O staining (Figure 6A). We explored whether release of growth factors after injury was blunted in OA and whether it was dependent upon severity of disease in an aggrecan-dependent manner. Cartilage biopsies were taken from human osteoarthritic cartilage obtained from knee arthroplasty surgery (Figure 6B) and scored for macroscopic disease severity using the Outerbridge classification. Proteoglycan content (as assessed by amount of GAG) inversely correlated with Outerbridge score (Figure 6C), confirming that severe macroscopic damage was associated with reduced GAG (aggrecan) content. Uniform biopsy punches (matched by wet weight) from severely diseased regions (Outerbridge score 3–4) released significantly less HDGF, CTGF and FGF2 upon injury compared with biopsies taken from relatively undamaged regions of the same joint (Outerbridge score 0—1) (Figure 6E–H). This was not due to reduced growth factor content as protein extracted from the matrix with 2.5M NaCl indicated that the more damaged cartilage contained higher Na-extractable growth factor compared with tissue with mild disease (Figure 6I–L).

**Fig. 6.**
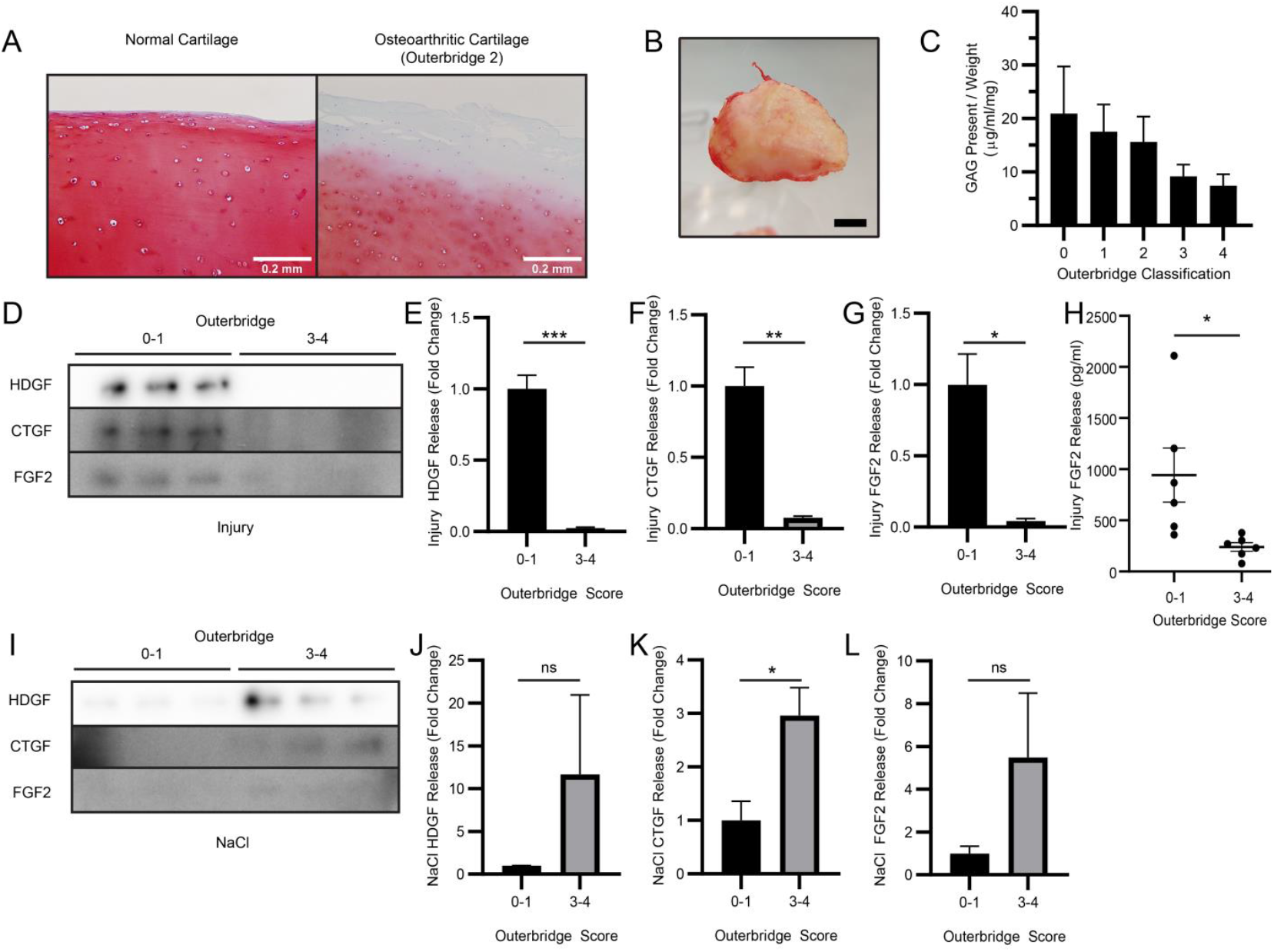
Release of heparan sulfate-bound growth factors is reduced from diseased osteoarthritic tissue compared to less damaged tissue after cutting injury. (A) Healthy and osteoarthritic human articular cartilage stained with Safranin O (0.2 mm scale). (B) Human osteochondral samples were collected from arthroplasty speciments and osteoarthritis severity scored macroscopically according to the Outerbridge Classification (1 cm scale). (C) Individual explants were digested with proteinase K and assessed for GAG content by the dimethylmethylene blue assay. (D-G) Injury conditioned medium (15 mins) was collected from osteoarthritic cartilage samples from mildly damaged (0-1) or severely diseased (3-4) areas and immunoblotted for growth factors, (H) or assessed by ELISA (FGF2). (E-G) Quantified from D (representative image, n=3 biological replicates). Injured cartilage explants were extracted with 2.5 M NaCl. (I) Matrix extract was immunoblotted for growth factor release. (J-L) Quantified from I (representative image, n=3 biological replicates). T-test (*=p<0.05, **=p<0.01, ***=p<0.001).

## Discussion

To date our group has identified three growth factors that are released rapidly following mechanical injury of articular cartilage ^3,13,20^. Here we characterise, for the first time, the release of HDGF, a lesser known heparan sulfate bound growth factor, identified in a proteomic analysis of articular cartilage PCM. Like CTGF and FGF2, HDGF was released rapidly, in a cell-independent manner, from a heparan sulfate bound matrix store. Commonality between the three growth factors, in terms of release by injury and sequestration on heparan sulfate within the PCM, suggested a common mechanism of release.

Three distinct properties of cartilage led us to hypothesise that release upon injury was due to a shift in the matrix sodium concentration. Firstly, the extracellular sodium content in cartilage is high compared with other tissues, at around 250-350mM ^24–27^, which we confirmed by ^23^Na MRI imaging (Figure 3B). Secondly, we identified a gradient of sodium between the territorial/interterritorial matrix and pericellular matrix due to absence of aggrecan (chondroitin sulfate) in the PCM (Figure 2A). Thirdly, all three growth factors were bound to heparan sulfate in cartilage (Figure 1F, G) and had previously been shown to elute from heparin using NaCl ^19,28^. Our previous studies had demonstrated that perlecan is the major heparan sulfate proteoglycan in cartilage and is exclusively found in the PCM ^3^. Though we were unable to immunolocalise HDGF in the PCM (data not shown), it was highly likely that it was also bound to perlecan along with CTGF and FGF2.

Our data support a sodium dependent release of growth factors upon injury by demonstrating that modulating sodium directly (by exogenous NaCl addition) or indirectly (by depleting aggrecan), increases or decreases, respectively, the release of growth factors after injury. Moreover, a shift in free sodium, relative to bound sodium, within the tissue was demonstrated after cartilage compression using ^23^Na MRI. This indicates that pathological type mechanical loading (28% unconfined strain) leads to a relative increase in free sodium levels and decrease in the bound sodium levels.

Regional changes in sodium observed by ^23^Na MRI suggested that sodium mobilisation occured non-uniformly in the cartilage upon compression. This was consistent with regional CTGF/TGFβ activation (Smad 2 phosphorylation), which occurred just below the articular surface upon loading, and differences in matrix stiffness. Indeed, this region of matrix had a different polarised microscopy appearance, consistent with the known random alignment of collagen fibres in this region ^29^, and displayed reduced tissue stiffness, which became more evident upon compression (Figure 4F). These observations were also confirmed in human cartilage, where a marked reduction in volume was observed in the area just below the superficial zone, but not the deep zone, which was associated with marked collagen fibre realignment (Figure 5). Our data are consistent with previous findings that the region just below the surface is lower in fixed charge density ^24,30^, has increased permeability ^31^, and is more likely to compress upon load ^30,32,33^. It is also where the tissue exhibits high cation fluxes ^34^.

Fluid flows away from more readily compressed regions to unloaded regions ^35^. Fluid flow is also high in osteoarthritic tissue which loses its fixed charge density, has a lower compressive modulus (more susceptible to compression) and becomes more permeable ^36–38^. However, fluid transport is not thought to significantly affect the transport rate of small solutes in cartilage, as diffusion coefficients are the dominant factor for small solute transport ^39^. Therefore, we suggest that displacement of growth factors upon loading occurs as a result of the net change in the local concentration of free sodium in the tissue rather than due to movement of sodium mediated by fluid flow. Taken together, we suggest that the region just below the articular surface selectively loses water, causing a local increase in the concentration of free sodium, and this drives growth factor release when compression is applied.

This mechanism fails in osteoarthritis where severe loss of proteoglycan leads to an inability to release growth factors upon injury. Growth factors are still present in the tissue and can be extracted by NaCl; indeed these appear to be increased in severe disease compared with less diseased osteoarthritic tissue. Proteoglycan loss and loss of fixed charge density are well established pathological features of osteoarthritis ^40,41^, and this is mediated by specific aggrecan degrading enzymes such as a disintegrin and metalloproteinase with thrombospondin motif-5 (ADAMTS5) ^22^. Aggrecanase activity is an early feature of disease and may precede collagen degradation ^42^. Early partial GAG loss, when the collagen network is still intact, can render cartilage more sensitive to load and increased fluid loss ^43^ suggesting that in early disease there may be enhanced release of growth factors. This could explain why observed release of growth factors was strong from the less affected osteoarthritic cartilage explants (Outerbridge score 0–1) (Figure 6D). We were unable to test whether this was higher than the amount released from completely normal articular cartilage, due to difficulties obtaining such tissue.

Enhancement of release early in disease might therefore represent a previously unrecognised role for aggrecanases in promoting the intrinsic repair response of articular cartilage. Once proteoglycan is severely depleted, osteoarthritic tissue loses this mechanism of growth factor release and this presumably then contributes to the failure of the tissue.

Loss of cartilage repair due to reduced growth factor bioavailability is emerging as an important mechanism in osteoarthritis pathogenesis. Both FGF2 and TGFβ have disease modifying roles in pre-clinical models *in vivo* ^11–13^. FGF2 signalling is known to enhance the chondrogenic potential of mesenchymal stem cells by priming them for chondrogenesis ^44–46^, and TGFβ drives chondrogenic differentiation ^14,15,47^. Both FGF and TGFβ pathways have been identified in osteoarthritis genome wide association studies, where hypomorphic variants are associated with increased risk of disease ^48^,^49^. Both growth factors, FGF2 and TGFβ, predict clinical response to surgical joint distraction in OA patients; a procedure that drives regeneration of articular cartilage ^50^. The therapeutic potential of intrinsic growth factors is reflected in the recent successful phase 2 FORWARD trial (NCT01919164) of a truncated form of recombinant fibroblast growth factor 18 (sprifermin). This study is the first structure modifying drug to demonstrate efficacy by building back articular cartilage in osteoarthritis ^51,52^. The role of HDGF is currently unknown, but it has been implicated in the tissue injury response in smooth muscle cells of vascular tissue ^53,54^. Its functional role *in vivo* is being explored.

Our data, supported by historical studies on tissue composition, response to mechanical load and solute transport, support a model whereby cartilage compression expels water preferentially from less stiff regions of the matrix, causes an increase in free sodium concentration, and enables the release of growth factors from the PCM. This study identifies an important intrinsic repair mechanism of normal cartilage that is lost in osteoarthritis, but could be exploited to recover reparative activities to improve disease outcome.

## Materials and Methods

### Tissue

Porcine articular cartilage was from the metacarpophalangeal joint for injury and recutting experiments, or from the medial tibial knee joint for imaging experiments, of 7-month-old pigs obtained from an abattoir after slaughter. Human articular cartilage was obtained from knee arthroplasty surgery of osteoarthritic patients.

### Cartilage Injury

For recutting injury experiments, uniform cartilage discs were explanted using a 4 mm biopsy punch (Stiefel #BI1500). Explants were rested in serum-free Dulbecco’s Modified Eagle’s medium (DMEM) for 48 h. Rested explants remained uncut (rested) or were injured (recut into 4 pieces by cutting twice with a scalpel (Swann-Morton, 15 blade) for 30 min, or specified time, in serum-free DMEM, after which time the conditioned medium was removed for immediate use or stored at −20 °C. Dead cartilage was freeze-thawed after dissection and washed thoroughly before recutting experiments. Cartilage injured at 4 °C or in the presence of inhibitors were pre-treated for 15 min prior to remaining uncut (rested) or being injury (recut), using batimastat from AbCam (ab142087) or protease inhibitor from Roche/Sigma (#4693132001). Recutting experiments in the presence of exogenous NaCl were conducted by placing explants in serum-free DMEM containing added NaCl to a total of (137 mM; 1x, 274 mM; 2x, 548 mM; 4x, 822 mM; 6x). After 15 min, conditioned medium was removed for immediate use or stored at −20 °C.

### Aggrecan depletion of chondroitin sulfate (sodium depletion)

Uniform cartilage discs were explanted using a 4 mm biopsy punch and rested in serum-free Dulbecco’s Modified Eagle’s medium (DMEM) for 24 h. Explants were treated either with serum-free DMEM or with serum-free DMEM containing 50 ng/ml interleukin-1 for 7 days. Conditioned medium was collected and replenished every 24 h, and assessed for GAG release by DMMB assay. After 7 days, recutting injury was performed as described, or explants were stained with safranin O to assess fixed charge density. Alternatively, after 7 days, explants were extracted using a dissociative whole tissue lysis buffer (RIPA), or matrix extraction buffer (2.5 M NaCl).

### Sodium ^23^MRI Loaded Imaging

Osteochondral cubes of 1 cm^3^ were dissected from the porcine knee medial tibial plateau, then vacuum sealed and placed in a custom-made compression cell. Imaging and spectroscopy were conducted for both proton and sodium nuclei on a Bruker 9.4T Ultra-high field microimaging system. Samples were imaged firstly uncompressed, then after 28% deformation compressive load to the cartilage surface, and finally after relaxation with no compression.

Loading in MRI experiments was performed after placing samples into a custom-made compression cell. The cartilage specimen was placed in the centre of the cell and compressed to approximately 30% of the original cartilage thickness using a screw-driven plunger entering the cell from the top. The compression was released by returning the screw-driven plunger into its original position. Samples were imaged firstly uncompressed, then after 28% deformation compressive load to the cartilage surface (as monitored by proton MRI), and finally after relaxation with no compression.

Anatomical images were acquired with home written gradient echo code (TopSpin3.2, Bruker, Germany) using dual tuned 1H/23Na microimaging coil to ensure proper co-registration of anatomical and sodium images. Each anatomical imaging protocol was performed without slice selection in order to match best sodium scan parameters. Time domain data were acquired into 256×128 matrices and were processed using Prospa 3.2 (Magritek, Germany) to result in the magnitude images with FOV = 19.52 mm2 and in plane image resolution of 76 μm2. Further experimental details are reported in the Supplementary.

Sodium imaging was performed after ^1^H scans. Localisation of free and bound sodium was performed with TopSpin 3.2 (Bruker, Germany) using a home written non-selective Gradient Echo (GE) MRI protocol. Bound sodium was visualised during loading cycle by inserting multiple quantum filter (MQF) into GE protocol before applying localising gradients. MQF time duration was determined from a series of triple quantum filtered experiments and set to the evolution time τ where the triple quantum filtered signal was at maximum. Achieved in plane resolution in sodium images was (117 × 0.109) μm^2^ for free and (62 × 86) μm^2^ for bound sodium. All details pertinent to sodium imaging including MQF imaging are given in the Supplementary.

Two dimensional (2D) ^23^Na MRS TQ TPPI spectroscopy was performed during loading cycles in order to determine changes in free and bound sodium levels upon compression and relaxation. All details of these experiments and analysis of the obtained spectra to yields fractions of sodium in the free and bound states during the loading cycle are reported in the Supplementary.

### Cartilage Depth Loaded Imaging

Porcine articular cartilage explants were taken from the pig knee joint by a 4 mm biopsy punch (Stiefel #BI1500). Explants were washed in serum-free DMEM and serum starved for 48 h. Cartilage explants were either cyclically loaded (0.1Hz) or rested in serum free DMEM for 45 min at 37°C and snap frozen in liquid nitrogen. Frozen explants were mounted in optimal cutting temperature compound (OCT) and cut by cryostat into 7 μm frozen sections (through the full thickness) on Surgipath snowcoat Xtra slides.

Alternatively, rested porcine knee joint cartilage samples were cut across specified depths for mass spectrometry analysis or scanning acoustic microscopy and polarised light microscopy.

### Sample preparation for mass spectrometry analysis

4M Gu-HCl tissue extracts of sectioned regions (from superficial zone towards the bone) were precipitated with ethanol (9:1) overnight at 4°C and collected after centrifugation at 13,200 g at 4°C for 30 minutes. The precipitate was dissolved in 100 μL of buffer (25 mM ammonium bicarbonate (AmBic), pH:7.8) /0,2% Rapigest (Waters) reduced with 2mM DTT at 56°C for 30 minutes on an orbital shaker and alkylated with 8 mM iodoacetamide at room temperature for 1h in the dark. Trypsin digestion was performed with 2μg of trypsin gold (Promega, Madison, WI) at 37°C on a shaker for 16h. Subsequently, samples were diluted to 200μl with 1M AmBic and filtered through a 30kDa filter (Pall Life Sciences, Port Washington, NY) by centrifugation at 2060 x g for 8 minutes. The filter was then washed with an additional 100μl 0.5M AmBic buffer to optimize recovery. The filtrates were then desalted using a reversed-phase C18 column (UltraMicro Spin C18, The Nest group, Southborough, MA) according to the manufacturer’s instructions.

### Discovery mass spectrometry

The samples were analyzed on a quadrupole Orbitrap benchtop mass spectrometer (Q Exactive™) (Thermo Fisher Scientific, Waltham, WA) equipped with an Easy nano-LC 1000 system (Thermo Scientific, Waltham MA). Separation was performed on 75 μm × 25 cm capillary columns (Acclaim Pepmap™ RSLC, C18, 2μm, 100Å, Thermo Scientific, Waltham, WA). A spray voltage of +2000 V was used with a heated ion transfer setting of 275°C for desolvation. On-line reversed-phase separation was performed using a flow rate of 300 nl/min and a linear binary gradient from 3% solvent B for 60 min to 35% solvent B, then to 90% solvent B for 5 min and finally isocratic 90% solvent B for 5 min. An MS scan (400-1200 m/z) was recorded in the Orbitrap mass analyzer set at a resolution of 70,000 at 200 m/z, 1×10E^6^ automatic gain control (AGC) target and 100 ms maximum ion injection time. The MS was followed by data-dependent collision induced dissociation MS/MS scans at a resolution of 15,000 on the 15 most intense multiply charged ions at 2 × 10E^4^ intensity threshold, 2 m/z isolation width and dynamic exclusion enabled for 30 seconds.

### Database search

Identification was performed using the sus scrofa taxonomy (22,166 sequences) setting of the Uniprot database (Proteome UP000008227) with Proteome Discoverer 2.5 (version 2.5.0.400,Thermo Scientific). The processing workflow consisted of the following nodes: Spectrum recalibration; Minora Feature detection; Spectrum Selector for spectra pre-processing (350-5000 Da); Spectrum Properties Filter (S/N Threshold: 1.5); Spectrum Properties (charges 2–6); Sequest-HT search engine (Protein Database: see above; Enzyme: Trypsin; Max. missed cleavage sites: 2; Peptide length range 6–144 amino acids; Precursor mass tolerance: 10 ppm; Fragment mass tolerance: 0.02 Da; Static modification: cysteine carbamidomethylation; Dynamic modification: methionine and proline oxidation; INFERYS Rescoring and Percolator for peptide validation (FDR<1% based on peptide q-value). Results were filtered to keep only the Master protein and protein FDR <1%. The quantification workflow was based on protein abundance using top-3 peptide intensities average allowing only unique peptides.

The mass spectrometry proteomics data have been deposited to the ProteomeXchange Consortium via the PRIDE ^55^ partner repository with the dataset identifier PXD023890.

### Immunohistochemistry

Frozen tissue sections were fixed in methanol for 10min, blocked with 10% goat serum/PBS for 60min at room temperature, and incubated with anti-phospho-Smad2 antibody from Cell Signaling (#138D4) (1:1000 dilution in goat serum) for 1hr at room temperature. Sections were washed three times with PBS and incubated with secondary Alexa 647-conjugated IgG, goat anti rabbit antibody from Invitrogen (#A-31852) for 1h at room temperature. After wash and mounting in 50% glycerol, florescence signals were detected by Ultraview confocal microscopy (Perkin-Elmer, x60 oil immersion lens).

### Protein Analysis

Growth factor proteins were analysed in conditioned medium and dissociative tissue extracts by Western blotting using anti-HDGF from AbCam (ab131064), anti-CTGF from Santa Cruz (sc-14939) and anti-FGF2 from PeproTech (#500-P18). FGF2 in condioned medium was also analysed by FGF2 V-PLEX ELISA from MSD (#K151MDD).

### Scanning Acoustic Microscopy Loaded Imaging

Loaded and rested porcine articular cartilage was sectioned to 5 microns thickness. Sections were air dried overnight onto a charged glass slide. Sections were rehydrated and imaged under scanning acoustic microscopy (SAM) to measures acoustic wave speed, positively correlated to tissue stiffness. Images were taken from the articular (STZ) to the deep zone in normal and damaged samples to assess wave speed across the tissue by sampling 5 boxed regions (50×50 pixels) every 200μm. The same tissue region was imaged by polarised light microscopy.

### Human Osteoarthritic Cartilage

Human osteoarthritic cartilage was macroscopically graded for severity of disease by modified Outerbridge classification (Grade 0; normal cartilage, Grade 1; cartilage with softening and swelling, Grade 2; partial-thickness defect with superficial fissuring, Grade 3; deep fissuring, Grade 4; exposed subchondral bone).

Human osteoarthritic articular cartilage biopsies were taken using a 4 mm biopsy punch, assigned to an Outerbridge classification. Explants were injured in 200 μl serum-free DMEM, after 30 min the conditioned medium was stored at −20 °C, then explants were fully digested with 50 μg/ml proteinase K and assessed for GAG content by DMMB assay. Alternatively, explant matrix proteins were extracted with 2.5 M NaCl for 30 min, then total tissue protein extracted with RIPA tissue dissociative lysis buffer for 1 h.

### Polarization sensitive SHG imaging of collagen fibres under compression

Polarization sensitive Second Harmonic Generation (PSHG) was used to investigate the collagen fibre realignments under compressive load on the microscopic scale. The strength of the SHG signal from collagen is highly dependent on the polarisation of the laser fundamental with respect to the collagen fibre axis. Therefore by taking a series of images at different polarization angles it is possible to extract detailed information on the collagen organisation within the tissue on the microscopic scale. This analysis provides two parameters, *I*_2_ and φ_2_, which describe the collagen fibre organisation on the microscopic scale. φ_2_ is the predominant collagen fibre orientation in a given pixel, and *I*_2_ is a measure of the molecular organisation at a submicron scale (*I*_2_=0 when the collagen fibre arrangement is isotropic, and increasing values of *I*_2_ indicate a more parallel collagen fibre arrangement). A full description of how the measurements are taken and how the parameters *I*_2_ and φ_2_ are derived is given in ^56^. 1 mm thick sections of human articular cartilage with the underlying bone attached were compressed on a bespoke loading rig which attached onto the stage of the multiphoton microscope (Supplementary Figure S3), after compressive loading the samples were left for 30 min to equilibrate.

## Supporting information

Supplementary Proteomics File

## Acknowledgements

This work was supported by funding from the Biotechnology and Biological Sciences Research Council (BBSRC) [grant number BB/M011224/1].

This work has been supported by the Centre for Osteoarthritis Pathogenesis Versus Arthritis (Grant 20205 and 21621).

Tissue samples and/or data were collected with informed donor consent in full compliance with national and institutional ethical requirements, the UK Human Tissue Act, and the Declaration of Helsinki (HTA Licence 12217 and REC 07/H0706/81).

The research was supported by the National Institute for Health Research (NIHR) Oxford Biomedical Research Centre (BRC). Disclaimer: The views expressed are those of the author(s) and not necessarily those of the NHS, the NIHR or the Department of Health.

Medical Research Council grant G1001398 [Quantifying age-related changes in the mechanical properties of tissues].

This work was partly funded by Arthritis Research UK grant No. 20714 and by a Leverhulme Trust Visiting Professorship (S. Brasselet) Grant No. VP2-2015-025.The collection and storage of human cartilage samples from total knee replacement surgery was carried out by the RD&E Tissue Bank, part of the NIHR Exeter Clinical Research Facility.

G.E.P thanks the Medical Research Council for funding (Grant No. MC_PC_15074) for developing sodium imaging methods.

This article contains supporting information online.

---

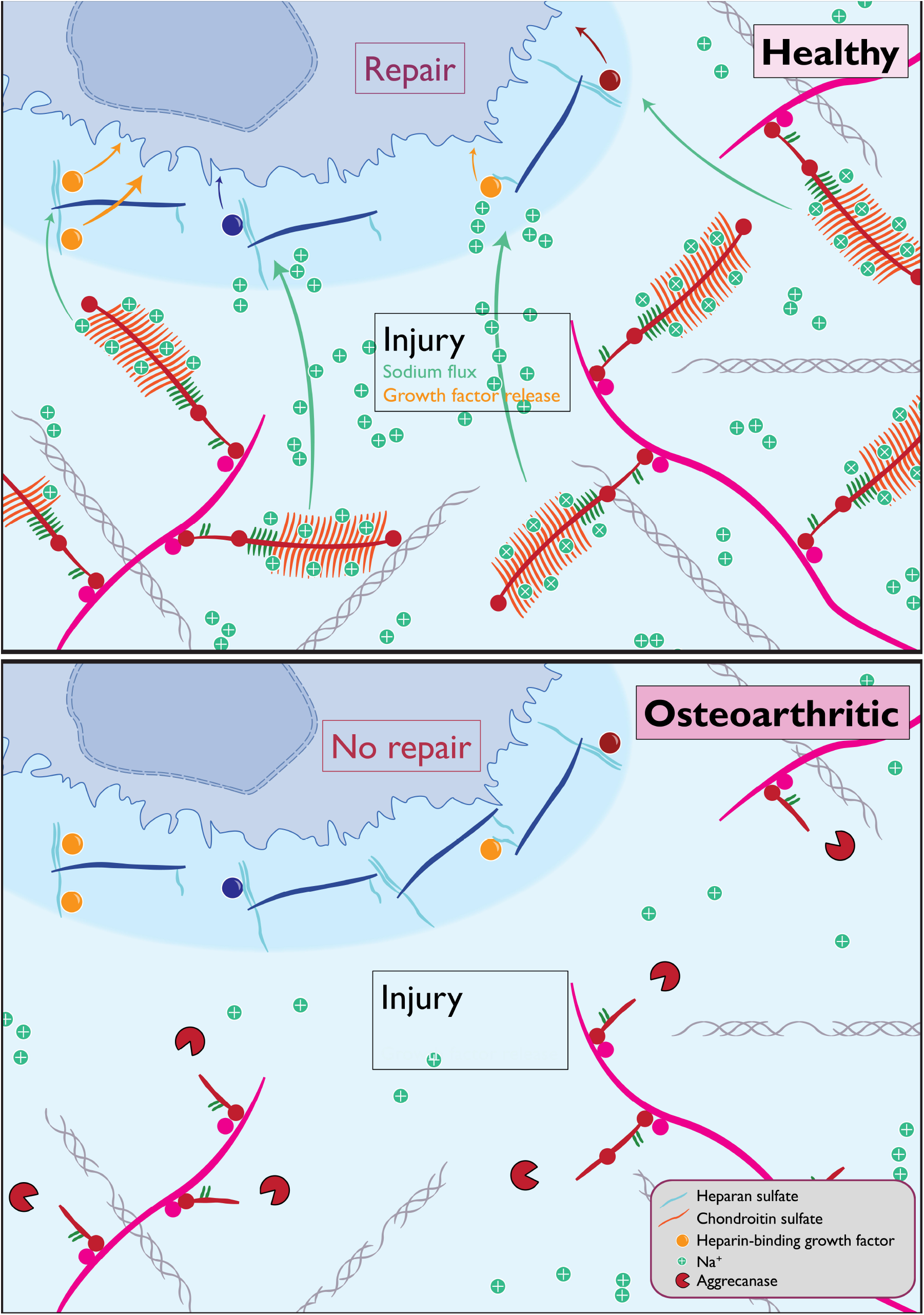

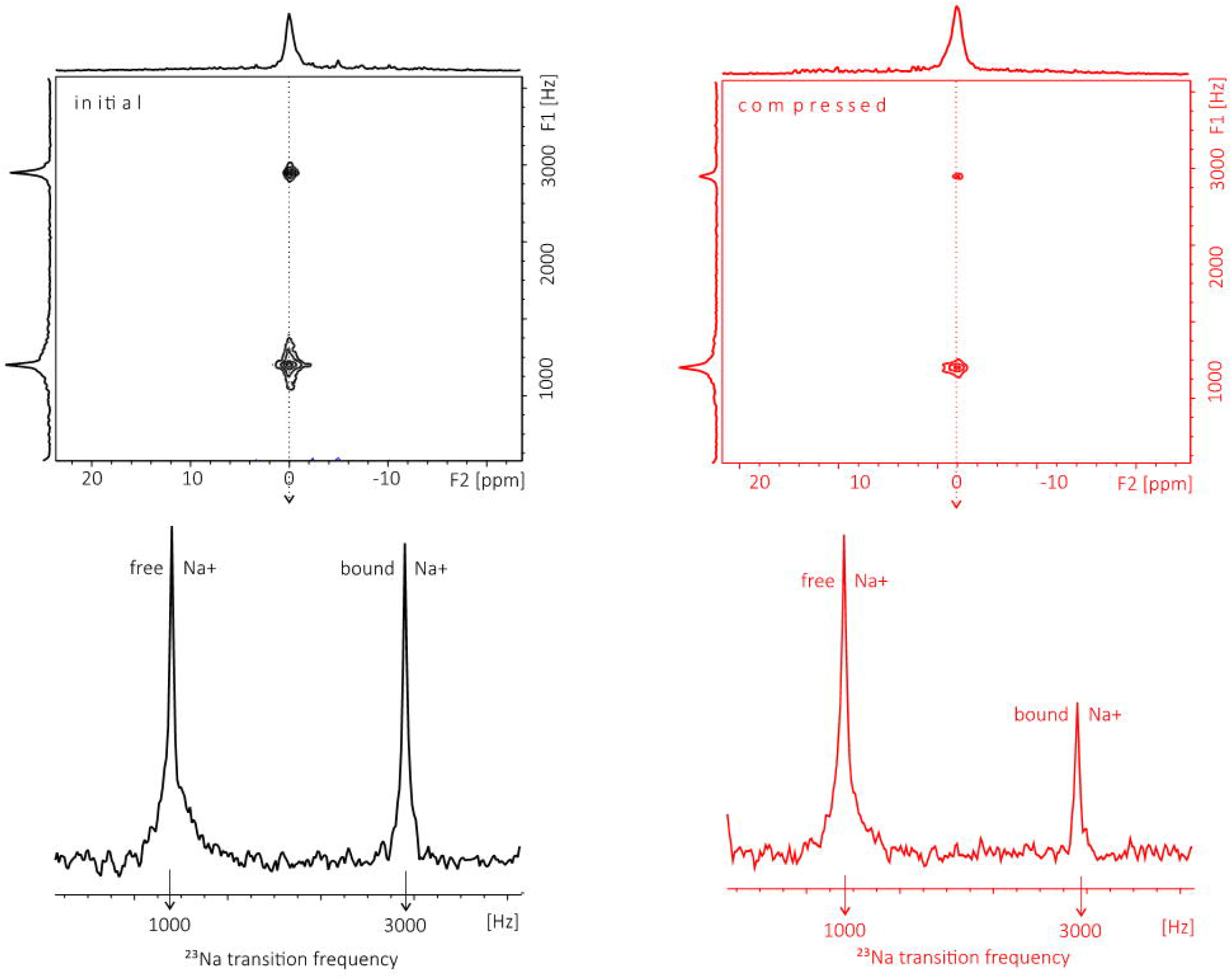

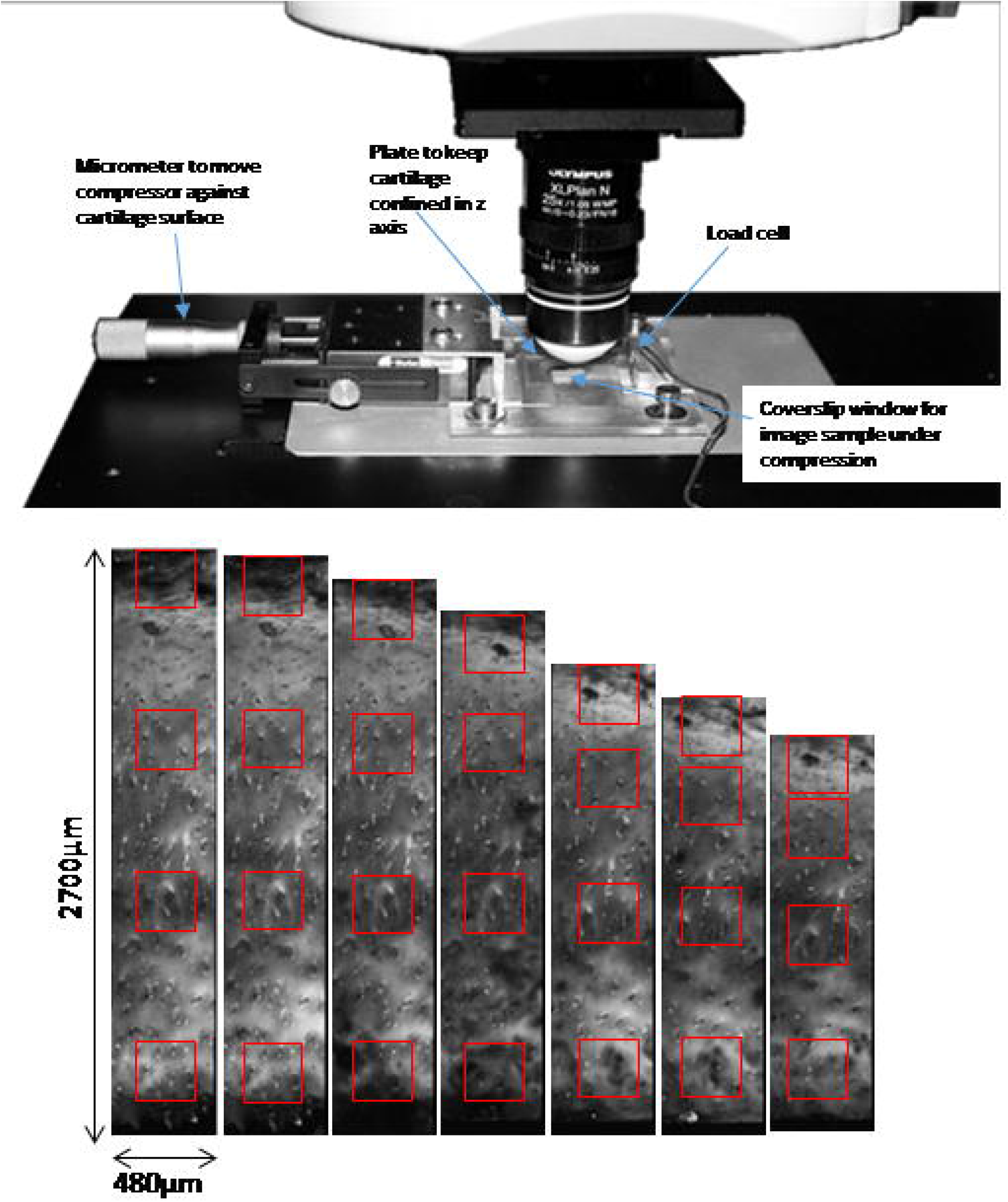

## Notes

http://proteomecentral.proteomexchange.org/cgi/GetDataset?ID=PXD023890

### Competing Interest Statement

The authors have declared no competing interest.

