## Supplementary for "Matrix-Bound Growth Factors are Released upon Cartilage Compression by an Aggrecan-Dependent Sodium Flux that is Lost in Osteoarthritis"

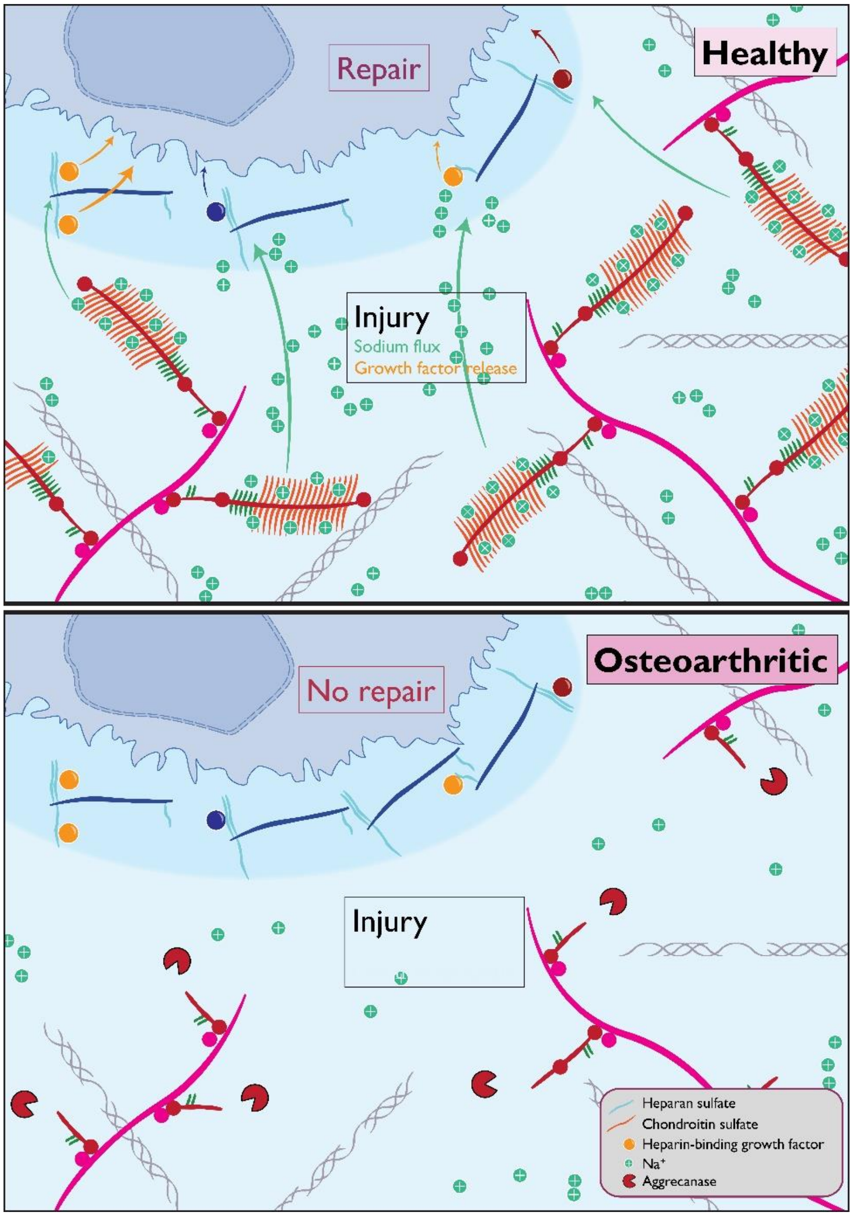


Figure S1. Schematic of aggrecan-dependent sodium flux upon cartilage injury in healthy and osteoarthritic tissue.

***MRI Sample preparation***

Osteochondral cubes 1 $cm^3$ in size were dissected from the porcine knee medial tibial plateau, then vacuum sealed and placed in a custom-made compression cell. One dimensional proton and sodium spectra of cartilage specimen used Imaging and spectroscopy were conducted for both proton and sodium nuclei on a Bruker 9.4T Ultra-high field microimaging system.

Compression/relaxation cycle was performed after placing samples into a custom-made compression cell. The cartilage specimen was placed in the centre of the cell and compressed to approximately 30% of the original cartilage thickness using a screw-driven plunger entering the cell from the top. The compression was released by returning the screw-driven plunger into its original position. Samples were imaged firstly uncompressed, then after 28% deformation compressive load to the cartilage surface (as monitored by proton MRI), and finally after relaxation with no compression.

### *Anatomical (1H) and Sodium (23Na) magnetic resonance imaging (MRI)*

NMR and MRI of all spin species reported in this study were performed on a 9.4 T Bruker Avance III Microimaging system (Bruker, Germany). Single-channel 25 mm $^{23}$Na and $^{1}$H commercial microimaging coils (Bruker, Germany), and a dual tuned $^{1}$H/$^{23}$Na microimaging coil (Bruker, Germany) were used in this study. Coils tuned to the resonance frequency of 400.18 MHz for protons and 105.86 MHz for sodium, were used for all $^{1}$H and $^{23}$Na porcine cartilage studies. The length of the $^{1}$H $\pi/2$ pulse was 32 μs at 150 W, the dwell time was set to 40 μs, and the resonance offset was set at the maximum of the proton peak height associated with water phase in each experiment or a series of consecutive experiments. One dimensional (1D) proton spectrum recorded for the cartilage specimen during compression/relaxation cycle did not reveal any significant changes in the spectrum indicating that no structural changes were induced by the loading cycle. The length of the $^{23}$Na $\pi/2$ hard pulse was 48 μs at 200 W, the dwell time was set to 80 μs, and the resonance offset was set at the maximum of the sodium peak height in each experiment or a series of consecutive experiments as only one peak was observed. 1D sodium spectra of a given cartilage specimen did not change during loading cycle also suggesting that structural integrity of the cartilage was preserved during the time-course of the loading cycle. All sodium experiments were performed at 37°C unless otherwise stated. Specific experimental spectroscopic and imaging details are provided below.

### *$^{1}$H MRI*

Anatomical images were acquired with home written gradient echo code (TopSpin3.2, Bruker, Germany) using dual tuned $^{1}$H/$^{23}$Na microimaging coil to ensure proper co-registration of anatomical and sodium images. Each anatomical imaging protocol was performed without slice selection in order to match best sodium scan parameters. Time domain data were acquired into 256x128 matrices and were processed using Prospa 3.2 (Magritek, Germany) to result in the magnitude images with FOV = 19.52 mm$^2$ and in plane image resolution of 76 μm$^2$.

### *$^{23}$Na MRS*

TopSpin 3.2 (Bruker, Germany) home-written $^{23}$Na MRS triple quantum filtered protocol (Jaccard, Wimperis and Bodenhausen, 1986; Navon *et al.*, 2001) was used to characterise time evolution of stored sodium in the skin. 23 triple quantum evolution delays $\tau$ were used to determine a delay $\tau_{max}$ where a maximum of triple quantum signal occurred. This $\tau_{max}$ delay was used in the design of the triple quantum filter used in the visualisation of bound sodium in the cartilage during the loading cycle. 960 transients were collected for each $\tau$ delay, with a recycle delay of 0.2 s, thus bringing the total time for the entire series to be within 2h.

Two-dimensional triple quantum-filtered with time proportional phase incrementation (TQ-TPPI) (Jaccard, Wimperis and Bodenhausen, 1986; Schepkin, 2016) incorporated into home written code for TopSpin 3.2 (Bruker, Germany) was used to determine fractions of sodium in free and bound states during compression/relaxation cycle. Spectral width in the direct dimension was set to 5 KHz, the indirect dimension was incremented using 512 50 μs time increments, recycle delay was 0.1 s, allowing for a spectrum collection in under 2 min. Data were processed with TopSpin3.2 with 20 Hz broadening in both dimensions and manual phasing of two-dimensional spectra in both dimensions. Fractions of free and bound sodium were produced by analysing central traces extracted at zero frequency in the direct dimension using build-in Multipeak fitting routine in IgorPro8.2 (Wavemetrics, USA). Integrated intensities of Lorentzian distributions obtained during spectral deconvolution were used as fractions of free and bound sodium observed in cartilage samples during loading cycle. Exmaples of 2D TQ TPPI maps and the traces extracted at zero frequency are given in Figure 1.

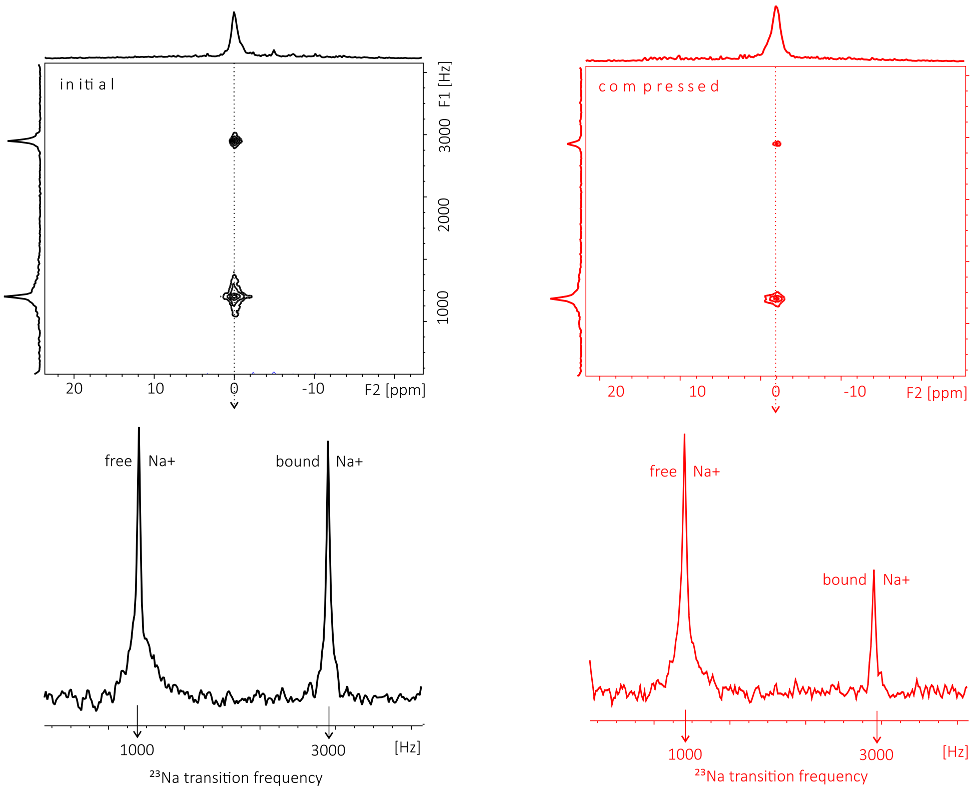


Figure S2. Two dimensional $^{23}Na$ TQTPPI maps of a cartilage specimen in the initial (black) and the compressed (red) states. One dimensional traces extracted at central frequency show initial sodium distribution in free and bound states (black) and change upon compression (red). Free and bound sodium fractions (n=4) shown in Figure 3C in the main text were determined from these traces using integrated intensities at each frequency determined by spectral deconvolution.

### ***$^{23}$Na MRI***

Localisation of free sodium was performed with TopSpin 3.2 (Bruker, Germany) using a home written non-selective Gradient Echo (GE) MRI protocol. The width of the $\pi/2$ pulse was 54 μs, the recycle delay was 0.2 s, the spectral width was 25 kHz. Images were collected into 64 × 64 matrices with the phase encoding time of 0.66 ms, and with maximum phase and read gradients set to 14% and 9%, respectively, resulting in a field of view, FOV = (30 × 28) mm$^2$. 512 transients were accumulated resulting in image collection within 1.8h. The images were reconstructed in Prospa 3.2 (Magritek, Germany) by zero filling to 256 × 256 matrices, sinc-squared apodisation performed in both dimensions with consecutive magnitude two-dimensional Fourier transformation. The plane resolution in processed images was approximately (0.117 × 0.109) mm$^2$.

Visualisation of bound sodium was performed using TopSpin 3.2 (Bruker, Germany) home written code where triple quantum filter was applied before localisation of bound sodium with the gradient echo. The evolution time in the filter was set to the $\tau_{max}$ value identified in the triple quantum filtered $^{23}Na$ MRS. The width of the $\pi/2$ pulse was 49 μs, with the width of $\pi$ pulse in the triple quantum filter set to 98 μs, the triple quantum filtered evolution time $\tau$ was set to 0.7ms, the recycle delay was 0.2 s, the spectral width was 50 kHz. Images were collected into 172 × 16 matrices with the phase encoding time of 0.32 ms, and with maximum phase and read gradients set to 9% and 13.8%, respectively, resulting in FOV = (16 × 22) mm$^2$. 960 transients were accumulated resulting in image collection under 1.8h. The images were reconstructed in Prospa 3.2 (Magritek, Germany) by zero filling to 256 × 256 matrices, sinc-squared apodisation performed in both dimensions with consecutive magnitude two-dimensional Fourier transformation. The plane resolution in processed images was (0.062 × 0.086) mm$^2$.

Sodium levels in studies cartilage samples were expressed in terms of signal-to-noise ratios (SNRs), to reflect relative sodium levels. Intensity images were converted to SNR images using the following expression (Wetterling *et al.*, 2012; Anisimov *et al.*, 2021):

$$SNR_j = \frac{S_j - \overline{N_S}}{\sigma(N_S)},$$

Where $S_j$ is the signal intensity measured in voxel $j$, $\overline{N_S}$ is the mean signal measured in the region of interest representing noise chosen as a square region with 6400 pixels from a region in the image containing no signal from the cartilage, and $\sigma(N_S)$ is the respective standard deviation of the noise mean. Multiple quantum filtering was utilised to distinguish between, visualise and characterise free and bound sodium fraction changes in the cartilage tissue during compression/relaxation cycle.

***Second Harmonic Generation Cartilage Loading Rig***

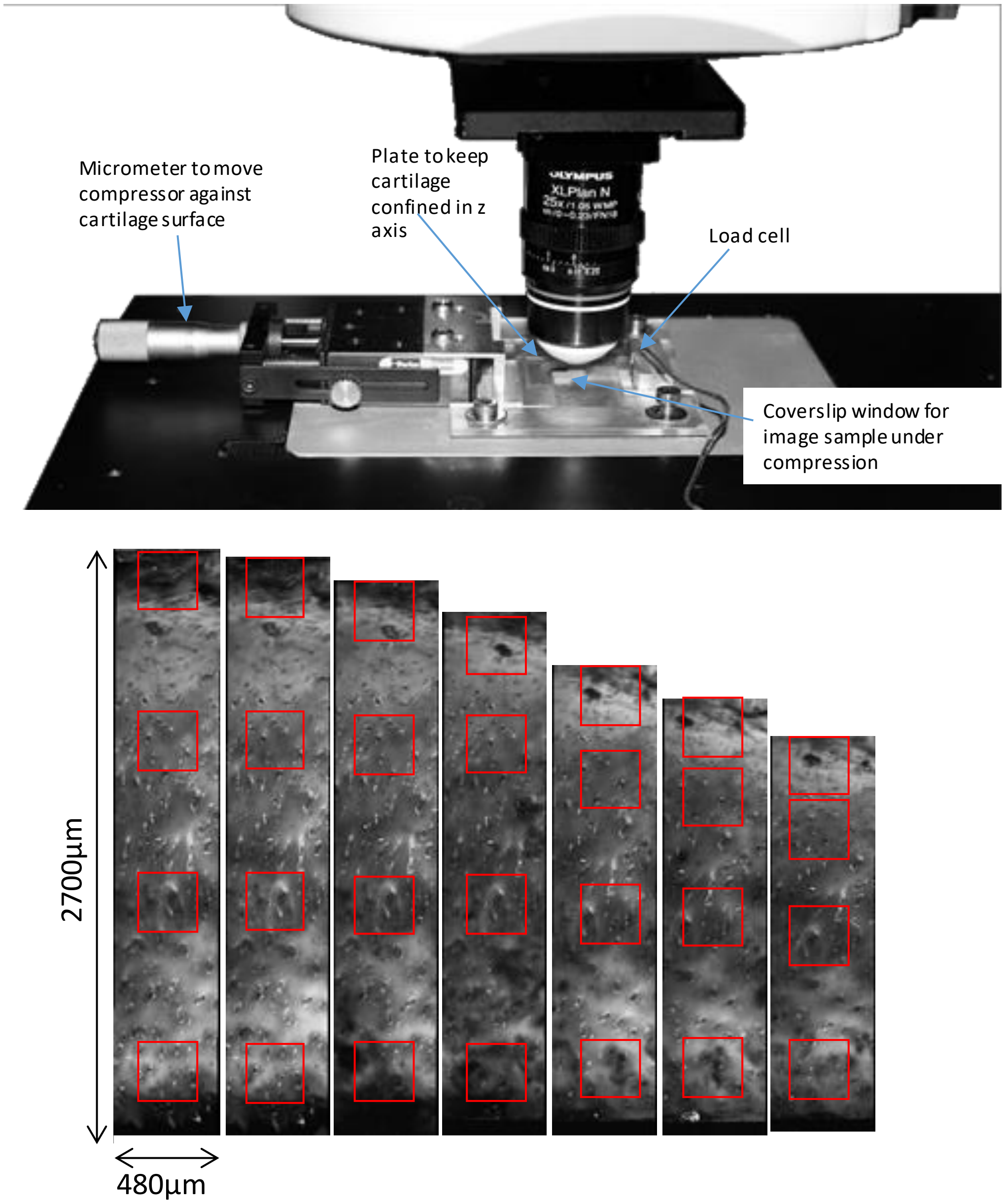


Figure S3. Second harmonic generation compression rig with articular cartilage sample from condyle of a 50-year-old male after total knee replacement surgery.
